# Swidden pollen spectra are unique, having no modern analogues

**DOI:** 10.1101/2025.07.14.664661

**Authors:** Ekaterina Ershova, Elena Ponomarenko, Valerii Pimenov

**Author notes:** Corresponding author: Ekaterina Ershova.

## Abstract

- Slash-and-burn cultivation (SABC), a widespread premodern agricultural practice, shaped vegetation across the temperate zone of Eurasia for millennia, yet the SABC-associated plant communities and their modern analogues remain poorly understood.
- We compared 74 fossil pollen spectra from radiocarbon-dated swidden horizons with 223 modern surface soil spectra using non-metric multidimensional scaling and cluster analysis.
- The results show that slash-and-burn cultivation creates a unique palynological assemblage in soil, distinguishable from forests, meadows, fallows, and ruderal habitats. This signature was most pronounced in sites from broad-leaved coniferous forest, where cryptogam diversity was highest.
- Here, we report that pollen spectra from buried swidden soils across the Eastern European forest zone consistently form a distinct and cohesive cluster, lacking modern analogues. These spectra are characterized by high proportions of *Betula, Tilia, Epilobium angustifolium*, cereal pollen, and a diverse assemblage of spores (*Marchantia, Lycopodium, Pteridium*), reflecting a mosaic of post-cultivation successional stages.
- Insect activity, particularly pollen-storing burrows of ground-nesting bees, may enhance the swidden signal through selective accumulation of fireweed entomophilous pollen.
- Our results provide SABC reference pollen spectra for temperate mixed and deciduous forest zones and can be used as a tool for reconstructing the extinct suite of plant communities associated with swidden cultivation.

## Introduction

Slash-and-burn cultivation (SABC), a variety of shifting cultivation that involves land clearance by fire, was practiced in temperate zones of Eurasia from the Bronze Age until the 1900-s (Conklin, 1957). Due to the short production stage and the long fallow stages of this agricultural technique, it affected plant communities of vast geographical areas, transforming their composition and, presumably, creating a suite of specific successional ecosystems. However, despite the ubiquity and obvious importance of this impact, it is unknown, what the plant communities associated with the slash-and-burn cultivation were and whether they have any analogues among modern ecosystems.

This problem can be approached by pollen analysis, one of the key methods for reconstructing past vegetation and land use history (Behre, 1986). It is most commonly applied to lake and peat deposits, where airborne pollen rain is well preserved and stratified. These aquatic archives provide regional-scale records of vegetation change, including signals of human activity (Alenius et al., 2022; Abraham et al., 2023).

In archaeology, the palynological analysis often complements reconstructions based on material finds, by documenting the environmental context of nearby sites. However, such archives are not always available, and their regional pollen signal may obscure short-term or small-scale episodes of land use—especially those involving insect- or self-pollinated crops or spatially localized features such as field boundaries (Bakels 2000; Hellman *et al*. 2009; Poska *et al.* 2018).

Soils, including those buried beneath archaeological layers or colluvial deposits, can also preserve pollen and spores—often of highly local origin. Pollen preservation in soils is more vulnerable to degradation due to aeration and biological activity. Unlike aquatic sediments, soils lack clear stratification, and a single horizon may contain pollen from multiple stages of vegetation development (Dimbleby, 1985). In addition to airborne pollen, soils accumulate pollen and spores from decaying plant material and may be further enriched by animals—for example, through herbivore manure or pollen stored in the burrows of ground-nesting bees (Bottema, 1975). As a result, soil and aquatic pollen spectrum of the same age and location may differ substantially (Lisitsyna *et al*., 2012; Ershova *et al*. in press), making interpretation more difficult. Nevertheless, we argue that soil pollen analysis can offer unique insights into short-term, small-scale, and site-specific characteristics, including land use practices.

The current approach to interpreting pollen data and past ecosystem dynamics relies largely on comparing subfossil pollen spectra with their modern or recent analogues. However, this method is not easily applicable to past anthropogenically modified ecosystems, which may themselves represent extinct landscape types with no counterparts in the modern world. Our experience shows that comparing pollen spectra from soils buried beneath archaeological layers with modern spectra—interpreted through the lens of present-day climatic analogues—can be misleading, often suggesting abrupt climate-driven ecosystem changes that are not supported by independent climate. Studies show that the variability observed in modern pollen spectra from the southern forest zone and forest-steppe is largely driven by land use rather than climatic gradients (Vyazov *et al*., 2019; Ershova *et al*. 2022; Lukanina & Shumilovskikh, 2025).

Ecosystems associated with “extinct” types of land use, such as slash-and burn (swidden) cultivation, are particularly elusive. While numerous archaeological cultures are believed to practice swidden agriculture at least from the Iron Age on, a distinct palynological signature of swidden cultivation in soil was described only recently by the authors (Ponomarenko *et al*, 2019). In this study, we aimed to answer the question whether pollen spectra of swidden horizons have any analogues among modern ecosystems of the temperate forest and forest-steppe zones [or swidden cultivation creates a unique combination of ecosystems that has to be treated as such].

## Objects and methods

### Buried SABC horizons

Over the past decade, we have studied traces of slash-and-burn agriculture in soils in the southern forest zone of Eastern Europe. A training set of swidden horizons was obtained in Karula NP (Estonia) based on historical evidence (Thomson *et al*., 2015), where diagnostic features of historical swidden horizons were described and first applied to identifying and radiocarbon dating ancient swidden horizons in the area (Ponomarenko *et al*., 2019). Further, we surveyed for similar horizons buried under various archaeological embankments (e.g., kurgans and fortification walls), and colluvial, aeolian, and fluvial deposits. Using this set of characteristics, we found and radiocarbon dated more than 120 buried slash-and-burn horizons in different regions of Eastern Europe (Ponomarenko *et al.*, in press).

The swidden horizons were identified based on their morphology and charcoal content. These are dark-colored layers, 4-10cm thick, with the lower boundary commonly dotted by round and tear-shaped infilled insect burrows, 1-1.5cm in diameter - trace fossils of sweat bees. The dark color is due too a high concentration of both macro- and microcharcoal, with 1 or more charcoal fragment larger than 2mm present in each cubic cm of soil mass.

Pollen analysis was performed on all swidden horizons using standard procedures for processing soil samples (Faegri & Iversen 1989). The most samples contained sufficient pollen for statistical analysis. In each region studied (Estonia, the Middle Volga region, the Moscow and Smolensk regions), we described a unique complex of pollen and spore taxa common to all swidden spectra in the region, distinguishing them from both surface spectra and other fossil spectra. In this work, we summarize the results of many years of palynological studies of swidden soils in order to identify regional and temporal differences, as well as common characteristics that could be used as universal indicators of ancient subsoil agriculture in ecological and archaeological studies.

For the analysis, we used data from previously published studies on Estonia (Ponomarenko *et al*., 2019), the Middle Volga region (Vyazov *et al*., 2019; 2021; Ponomarenko *et al*., 2020), Moscow (Ponomarenko *et al*., 2021; Ershova *et al*. in press), Smolensk (Ershova, Krenke 2017; 2025; Ershova, 2018; Ershova *et al.,* 2020; Krenke *et al*., 2019; 2021; 2022), and Kursk (Rodinkova *et al*., 2024) regions, as well as some unpublished data from archaeological reports (Supplementary Dataset S1). We selected only spectra from well-defined buried SABC horizons that preserved the entire complex of indicator features and were dated by radiocarbon and/or archaeologically. We excluded from the analysis samples that showed signs of redeposition, pronounced selective pollen decomposition (extremely low concentration and taxonomic poverty), as well as signs of mixing of several different stages of land use within a horizon (e.g., plowland after SABC and pasture after SABC). In total, we used 74 swidden spectra from 24 stratigraphic sequences located in the southern part of the forest zone and forest-steppe zone, of which 23 belong to the upper Dnieper basin, 15 to the Oka basin, 31 to the Volga basin, and 5 to the Baltic Sea basin (Fig. 1, Supplementary Dataset S1). The age of the SABC horizons ranges from ca. 4300 to 150 years ago (Ponomarenko *et al.,* in press).

**Fig. 1.**
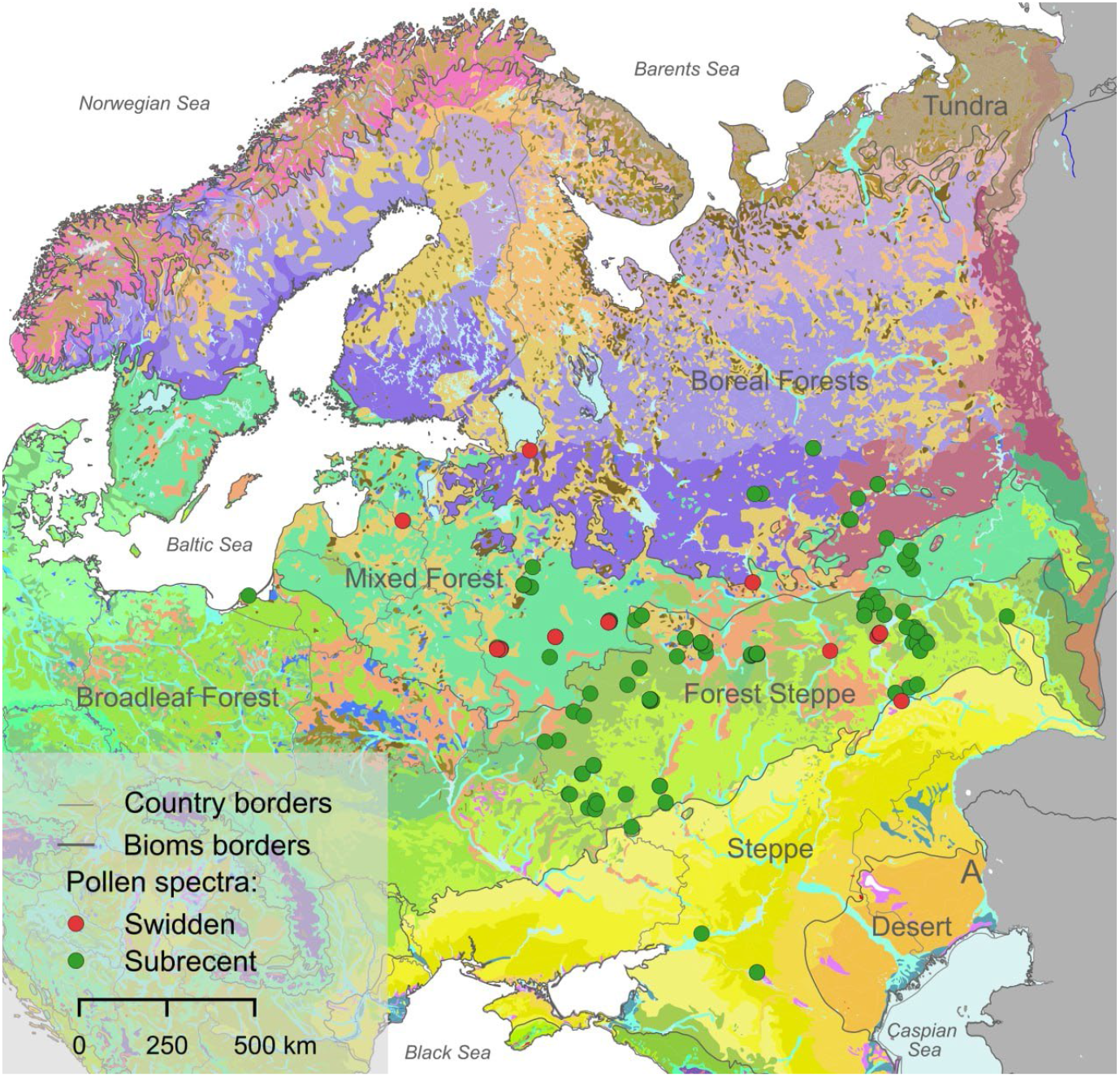
Map showing the locations of study sites overlaid on the Vegetation Map of Europe (after Bohn et al., 2004). Subrecent and swidden spectra are indicated by green and red circles, respectively. Biome boundaries are shown as grey lines.

### Surface soil spectra

To compare pollen spectra from buried SABC horizons with modern ones, we used published databases of surface spectra and our own data. Since pollen accumulates and preserves differently in soils than in aquatic deposits, we only used samples collected from the surface of mineral soils for comparison. 103 spectra with sample type “soil” and sample method “hand picking” were selected from the EMPD2 (Davis *et al*., 2020). The geographical distribution of the selected points is from 20 to 55 degrees longitude and from 46.5 to 60 degrees latitude, i.e., within the subzones of mixed coniferous-broadleaf forests, broadleaf forests, and forest-steppes. In addition, we used the results of our own studies of surface spectra (Ershova *et al*., 2022; Bakumenko & Ershova, 2021; Pimenov *et al*., 2025) and some unpublished data (Fig. 1; Supplementary Dataset S1). Surface spectra represent a wide range of local plant communities, both close to natural zonal communities and various agro-landscapes. A total of 130 surface samples represents forest communities, 54 represent meadows and pastures, 10 represent preserved steppe areas in the forest-steppe zone, 3 represent modern plowlands, and 25 represent abandoned plowlands at various stages of overgrowth (fallow). We also included various ruderal communities (roadside areas, wastelands, cattle tracks) in this group. For all pollen spectra from the surface of modern mineral soils, we use the term “subrecent,” since even when samples are taken from the top 1-2 cm, they contain pollen from more than one season.

### Statistical analysis

All pollen and spore taxa were used for the analysis. Surface and fossil soil spectra were taxonomically harmonized, original low-level taxa were grouped into higher-level taxa, and a total of 57 taxa were used. The proportion of each pollen taxon was calculated as a percentage of the total pollen, and the proportion of each spore taxon was calculated as a percentage of the total pollen and spores.

The TILIA and CONISS programs (Grimm, 1987) were used for unconstrained cluster analysis and diagram construction. Community patterns were investigated using non-metric multidimensional scaling (nMDS) using the vegan package (Oksanen et al., 2016) in R version 4.4 (R Core Team, 2024). The ordination was done using Bray-Curtis similarities of fourth-root transformed abundance data. The species significantly (p < 0.05) driving the separation of axes were visualised as biplots via the function envfit.

## Results and discussion

Figure 2 shows 222 surface and 74 fossil soil pollen spectra as a simplified pollen diagram constructed using unconstrained cluster analysis; the complete diagram is provided in Supplement Fig. S1.

**Fig. 2.**
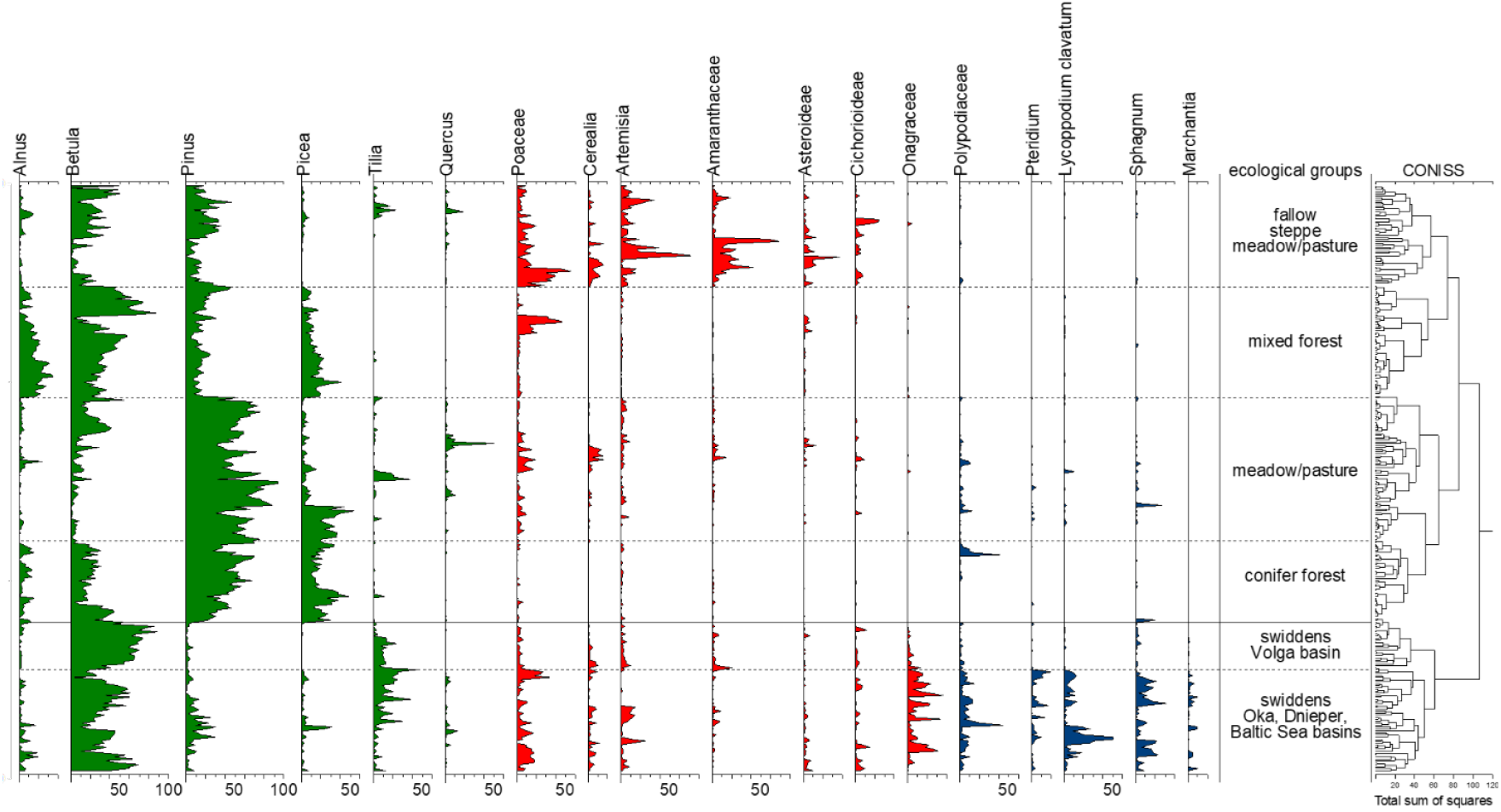
Surface soil pollen spectra and spectra from buried swidden horizons in the southern forest zone and forest-steppe of Eastern Europe. The spectra are grouped according to results of CONISS unconstrained cluster analysis. Pollen percentages are calculated relative to the total number of pollen grains; spores are expressed as a percentage of the sum of pollen and spores.

Fossil SABC spectra form a separate cluster, differing from all surface spectra in a number of parameters:

1. The dominance of *Betula* (up to 75% of the total pollen), often in combination with *Tilia* and a low proportion or absence of conifers.
2. The presence and often dominance (up to 40%) of Onagraceae (fireweed) pollen, a typical post-fire plant of the forest zone.
3. The abundance and taxonomic diversity of spores, especially *Lycopodium, Pteridium, Marchantia* and *Sphagnum*, which are indicator taxa for forest clearings and burnt areas.
4. The presence of pollen of cultivated cereals and some ruderal taxa (*Artemisia*, Amaranthaceae, Cichorioideae), which distinguishes SABC spectra from modern forest spectra.
5. A low proportion or even absence of meadow taxa, which, in combination with the dominance of birch, distinguishes swidden spectra from those of modern meadow and ruderal communities.

The presence of spores is typical for all analyzed SABC spectra, but two groups clearly stand out by their abundance (Fig. 2). In buried subsoils from the mixed coniferous-broadleaf forest subzone (Baltic Sea, upper Dnieper, and Oka basins), the number of spores is exceptionally high, accounting for up to 50% or more of the total. In buried SABC horizons of the Middle Volga region (broad-leaved forest and forest-steppe subzones), spores are present in much smaller proportions. This difference is obviously related to the much greater distribution of ferns, lycopods, and bryophytes in coniferous forests, where they are a very important component of the early successional stages following disturbances of the tree canopy and soil cover.

Figure 3 shows the result of non-metric multidimensional scaling (nMDS), which visualizes structural differences between pollen spectra from various modern ecosystem types and buried soils. Axes nMDS1 and nMDS2 represent a two-dimensional projection of multidimensional dissimilarities between spectra, based on their taxonomic composition (Bray-Curtis distances). Each point represents an individual pollen spectrum, point colors indicate ecosystem types. Blue arrows show taxon vectors, representing taxa that significantly contribute to axis separation.

**Fig. 3.**
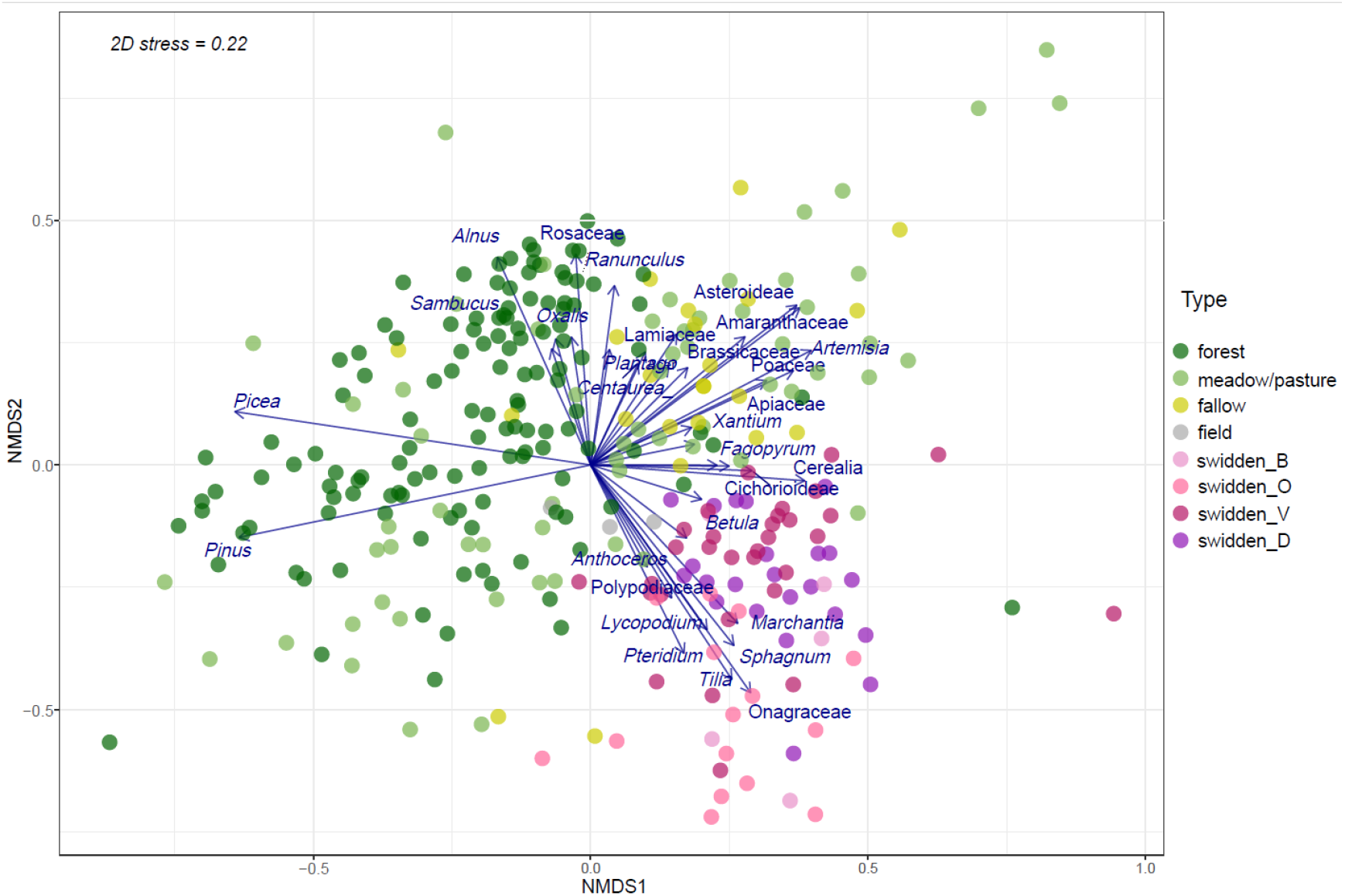
Results of non-metric multidimensional scaling (nMDS) analysis of fourth-root transformed pollen abundance data. Blue arrows indicate species significantly (p < 0.05) correlated with the ordination. The swidden spectra are colored according to their location in the basins of the Baltic Sea (swidden_B), Dnieper (swidden_D), Oka (swidden_O), and Volga (swidden_V).

Most forest spectra (dark green) are clustered in the left part of the plot, associated with *Picea, Pinus, Alnus*, and *Corylus*—taxa typical of the coniferous-broadleaf forest subzone and producing large amounts of pollen. The spectra of grasslands (meadows, pastures, hayfields) of the southern taiga subzone are located here as well, mainly in the lower left part (light green points).

The upper right part of the plot shows spectra dominated by herbs (Asteraceae, Amaranthaceae, *Artemisia,* Poaceae, Brassicaceae); these are mainly treeless areas of broad-leaved forests and forest-steppes subzones, as well as the vast majority of ruderal communities or fallows (yellow-green points).

All spectra of buried swidden horizons (pink and purple points) are concentrated in the lower right part of the plot, forming a separate compact group. Despite significant geographical and age variation, all studied buried SABC soil horizons contained a very similar complex of pollen and spore taxa: *Betula, Tilia*, Onagraceae, Cerealia, Cichorioideae, *Pteridium, Marchantia, Lycopodium clavatum, Sphagnum*, Polypodiaceae. This combination has no analogues among modern surface spectra of the entire southern forest and forest-steppe zones, despite a considerable diversity of their ecosystems.

The two spectra of young birch forests formed on former arable land and three spectra of modern plowlands were closest to the swidden spectra among the surface spectra. However, they lacked fireweed and spore-bearing plants, which are important indicators of swidden cultivation.

Modern fallow spectra (light yellow) are scattered on the nMDS plot (Fig. 3), but tend to cluster in the upper-right quadrant, overlapping partially with grassland and field spectra but clearly separated from the compact group of swidden horizons in the lower-right quadrant. They are associated with ruderal taxa such as *Artemisia, Plantago*, Cichorioideae, and Amaranthaceae, reflecting vegetation typical of abandoned or regenerating fields. In contrast to swidden spectra, fallow samples lack key indicators such as *Betula, Tilia, Epilobium* (Onagraceae), and a high abundance of cryptogam spores. This distinction suggests that modern fallows do not serve as analogues for swidden-related vegetation and supports the interpretation of swidden landscapes as a unique, now-extinct anthropogenic ecosystem.

## Conclusions

Buried soils that have been affected by slash-and-burn cultivation in the past contain a unique complex of pollen and spores that has no analogues among modern surface spectra in the southern forest zone of Eastern Europe. These spectra reflect a unique, now lost landscape— a mosaic of abandoned slash-and-burn plots at various stages of overgrowth. In addition to their specific morphology and charcoal pool, slash-and-burn horizons preserve a set of pollen and spores, the uniqueness of which can be explained by the combination of indicators of several stages of the slash-and-burn cycle: a production stage (cultivated cereals), initial stages of overgrowth (fireweed, bracken, liverworts, and lycopods), and advanced stages of overgrowth (young birch thickets). The unusual abundance of insect-pollinated fireweed pollen is probably associated with another characteristic of swidden horizons—signs of the activity of sweat bees, which could accumulate fireweed pollen in their burrows, thus giving the pollen spectra of SABC horizons their distinctive character.

Results of this study provide a novel tool for reconstructing the extinct suite of ecosystems/plant communities associated with the swidden cultivation and its past dynamics in space and time.

## Acknowledgements

This work was supported by the AAUW under American Short-Term Research Publication Grants [number 018870].

## Author contributions

Conceptualization (EE, EP), Data Curation (EE), Investigation (EE, EP), Writing — Original Draft (EE), Writing — Review & Editing (EE, EP, VP), Visualization (EE, VP), Funding acquisition (EE).

## Data availability

Data will be made available on request.

## Supporting Information

DataseSt1, DatasetS2, Fig.S1.

