## Supplementary figures and images for "Swidden pollen spectra are unique, having no modern analogues"

### Fig.S2.tif

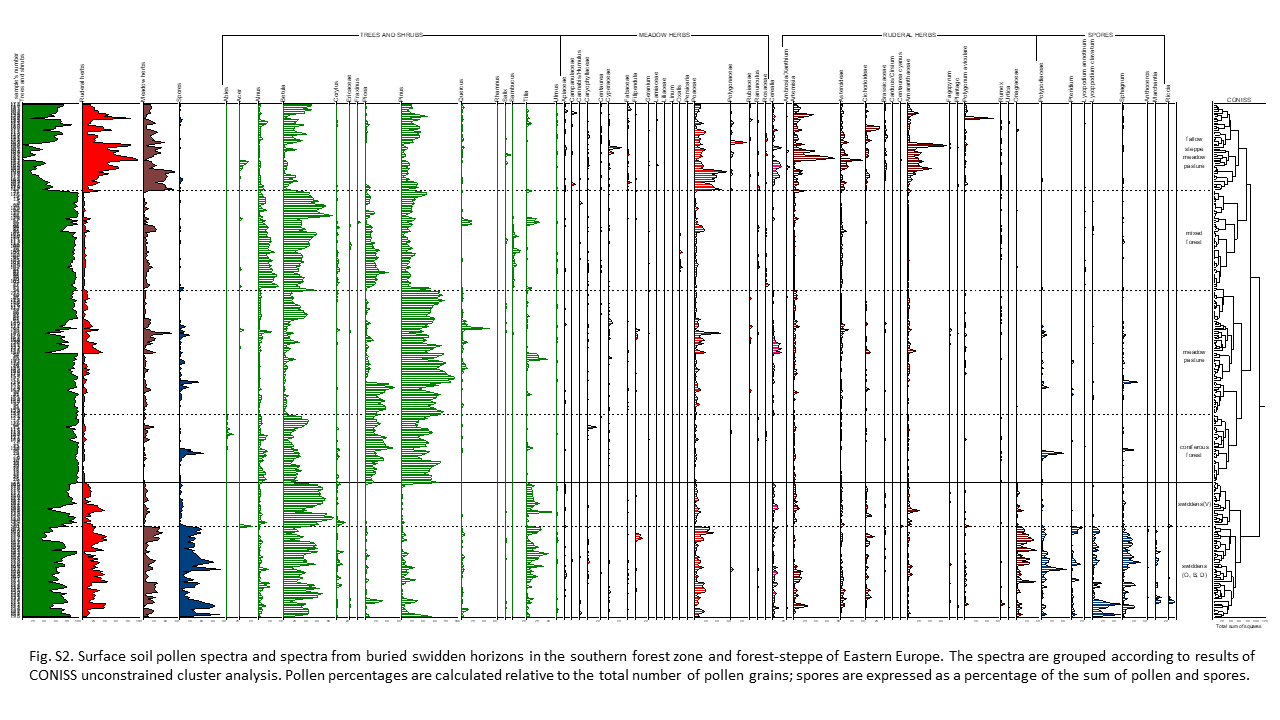
